# Genomic diversity in *Paenibacillus polymyxa*: Unveiling distinct species groups and functional variability

**DOI:** 10.1101/2024.03.26.586750

**Authors:** Adrian Wallner, Livio Antonielli, Ouiza Mesguida, Patrice Rey, Stéphane Compant

## Abstract

*Paenibacillus polymyxa* is a bacterial species of high interest, as suggested by the increased number of publications on its functions in the past years. Accordingly, the number of described strains and sequenced genomes is also on the rise. While functional diversity was repeatedly suggested for this species, the available genomic data is now sufficient to confidently segregate strains affiliated to *P. polymyxa* into two major species groups. As these strains exhibit considerable potential in agriculture, medicine, and bioremediation, it is preferable to clarify their taxonomic affiliation to facilitate reliable and durable approval as active ingredients. Strains currently affiliated to *P. polymyxa* can be separated into two major species groups with differential potential in nitrogen fixation, plant interaction, secondary metabolism, and antimicrobial resistance, as inferred from genomic data.

## Introduction

*Paenibacillus polymyxa* is a frequent inhabitant of various niches including soil, plant rhizospheres and plant tissues but also digestive tracts of different animals (Padda et al., 2017; Pasari et al., 2019). It has further been occasionally found in samples of seawater and fermented foods, but is also an inhabitant of the International Space Station (Daudu et al., 2020; Huang & Yousef, 2012; Langendries & Goormachtig, 2021). The species has undergone repeated taxonomic shifts and was reclassified as the type species of the novel genus *Paenibacillus* in 1994 (Ash et al., 1994; Judicial Commission of the International Committee on Systematics of Prokaryotes, 2005).

Due to its prolific secondary metabolism, manifold plant growth promoting potentials, probiotic status and bioremediation activities, *P. polymyxa* has been described as a model organism for sustainable plant, human and animal health management within the One Health framework (Pandey et al., 2023).

Strains of *P. polymyxa* are reported to provide multiple benefits for plant health in the shape of direct growth promotion. They increases nutrient availability through the conversion of atmospheric nitrogen to plant available ammonia, phosphate solubilization and iron scavenging (Puri et al., 2016; Raza & Shen, 2010; Xie et al., 2016; C. Zhou et al., 2016). *P. polymyxa* is also used as a biocontrol agent, known for its ability to produce volatile compounds that inhibit fungal plant pathogens (Lee et al., 2012; Rybakova et al., 2017). For example, the production of antifungal compounds by *P. polymyxa* is highly effective against *Fusarium* species, and this bacterium can also induce systemic resistance to pathogens in plants. (Beatty & Jensen, 2002; Dall’Agnol et al., 2017; Li & Chen, 2019). The overall beneficial traits of *P. polymyxa*, i.e. its potential for biofertilization, biocontrol and protection against abiotic stresses, have been recently reviewed by Langendries & Goormachtig (2021).

High interest in this species and simplified application of sequencing technologies have led to a leap in genome sequence depositions on public repositories. The first comparative genomics analysis for *P. polymyxa* is a decade old and included only four strains (Eastman et al., 2014). The pangenome of 14 strains was explored more recently (L. Zhou et al., 2020). In 2024, about 160 strains are affiliated to *P. polymyxa* when cross-referencing NCBI and GTDB databases. This considerable amount of genomic information coupled with the latest analysis tools enable a robust comprehension of this high-interest species.

Here we provide a species-wide comparative genomics analysis underlining the molecular oppositions between the different species groups currently affiliated to *P. polymyxa*. We also describe the putative functionalities of these groups on a basis of plant growth promotion traits, antimicrobial resistance, and secondary metabolism to obtain a comprehensive picture of their host interaction capacities.

## Results

### Strains affiliated to *P. polymyxa* segregate into distinct species groups

Assemblies for 110 *P. polymyxa* genomes are available in the NCBI RefSeq database. Their affiliation to *P. polymyxa* was confirmed using the GTDB toolkit (Chaumeil et al., 2022). The GTDB references 47 additional genomes belonging to *P. polymyxa* which in NCBI are either affiliated to alternative or unnamed *Paenibacillus* species. A total of 157 whole genome assemblies, representing 152 different strains are compared here. We kept the multiple entries from the same strains in our comparison as they have different genome assemblies.

Average Nucleotide Identity (ANI) is a whole-genome similarity metric allowing robust comparisons of taxonomic identities in prokaryotes (Goris et al., 2007). The ANI analyses conducted in a previous pangenome approach including 14 *P. polymyxa* strains concluded the existence of three species candidates (L. Zhou et al., 2020). Seven were considered part of the “true” *P. polymyxa* while six were attributed to *Paenibacillus peoriae* and one to a new species. All *P. polymyxa* strains included in the present analysis are genetically distinct from the *P. peoriae* type strain, as well as from any other type-strain of the 74 most closely affiliated *Paenibacillus* species (**Figure S1**).

Strains affiliated to *P. polymyxa* are separated in two major ANI groups, within which further subgroups can be distinguished (**Figure 1A**). All species groups have an average ANIb identity score below 90% similarity while the subgroups share an average internal similarity of at least 97.5% identity (Supplementary data). The cluster containing the *P. polymyxa* type-strain ATCC 842 will be further called species group A. The second dominant group will be referred to as species group B. A single strain, currently affiliated to *P. polymyxa*, can be clearly distinguished as an outgroup showing no significant similarity to any other strain within the *P. polymyxa* group nor to any other *Paenibacillus* type strain and will further be called species group C (**Figure 1A & Figure S1**). All alignments have a sufficient coverage to be interpreted as reliable results (**Figure S2**). The average whole-genome similarity between species groups A and B is 89,5%, considerably lower than the gold standard of 95% in vigor for most species delimitations (Richter & Rosselló-Móra, 2009). Based on nucleotide identity, species group B is closest to *Paenibacillus ottowii* (mean ANIm=89,3%) and to *P. polymyxa* species group A (mean ANIm=90.3%). Species group C is closest to *Paenibacillus xylanexedens* (ANIm=94.4%). We will further refer to the strains used in this analysis as the *P. polymyxa* group as they are currently affiliated to this species, even though ANI levels compared to other *Paenibacillus* species suggest that this group is not monophyletic (**Figure S1**). A phylogenomic reconstruction including five *P. ottowii* strains confirms that *P. polymyxa* species group A shares a more recent common ancestor with *P. ottowii* than with *P. polymyxa* species group B (**Figure S3**). Genomes identified as *P. polymyxa* only on GTDB are distributed in both major species groups (21 in species group A and 26 in species group B).

**Figure 1.**
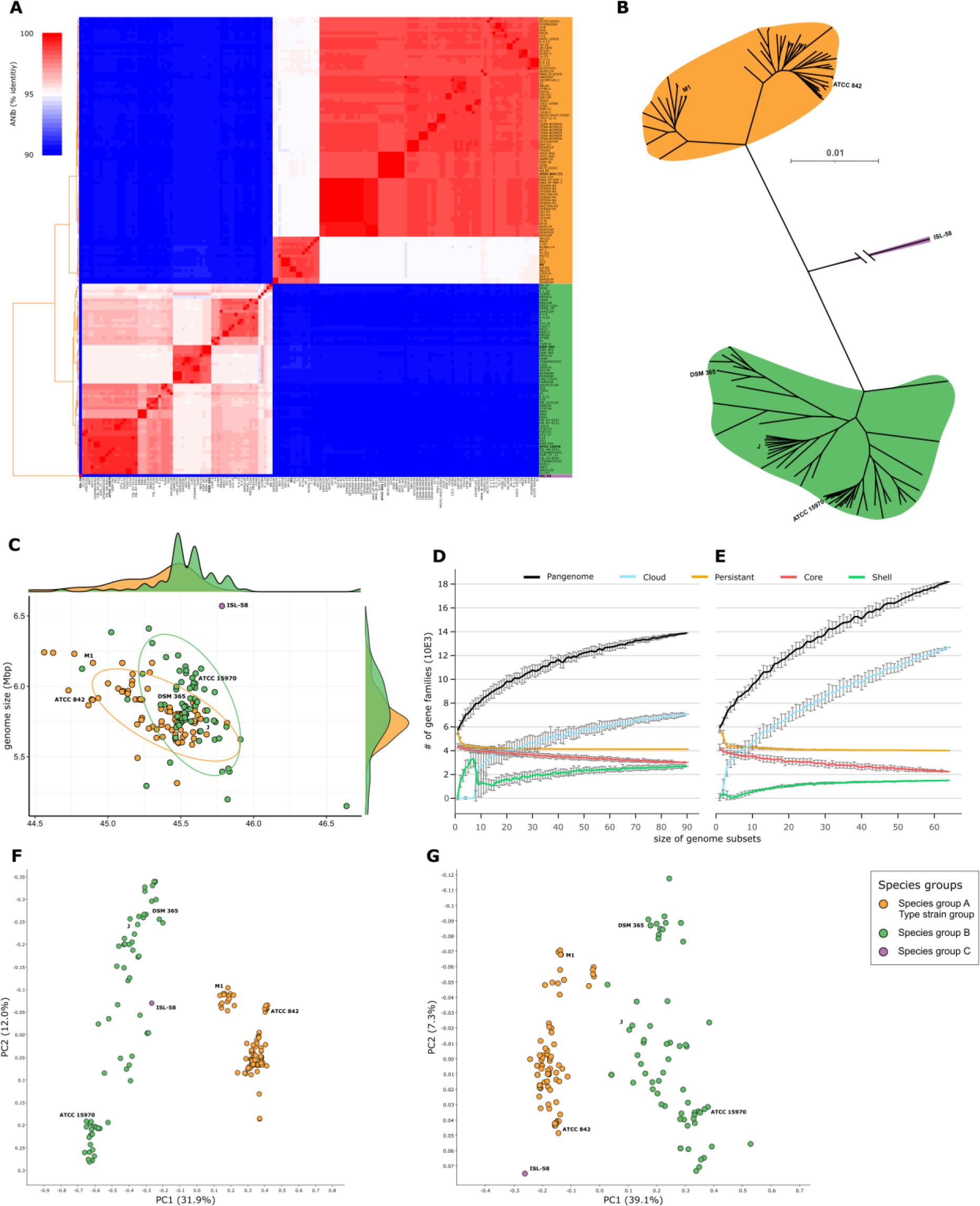
Segregation of *Paenibacillus polymyxa* strains. (A) Average nucleotide identity analysis based on pairwise blastn comparison of whole genome sequences between *P. polymyxa* strains (n=157). (B) Phylogenetic reconstruction of *P. polymyxa* affiliated strains. Single copy orthologous protein sequences were detected and aligned using Orthofinder resulting in an alignment of 1,397 sequences for a total of 406,116 amino acids. The tree was build using FastTree with default parameters. A rooted tree with local support values and complete leaf labels is available as supplemental data. Branch length for ISL-58 is 0.43 (C) Genome size and GC content comparison between *P. polymyxa* strains (n=157). The samples were colored according to the species groups determined through ANIb. (D) Rarefaction plots of pangenome calculations for species group A (E) and species group B. The different gene categories vary based on their frequency in the total observed population with cloud genes present in at least one organism, shell genes present in more than 8%, persistent genes present in more than 90% and core genes present in 100%. (F) Bray-Curtis PCoA representation of the genetic profile of *P. polymyxa* affiliated strains based on their pgpt-db affiliation or (G) their SEED affiliation.

A phylogenetic reconstruction was modelized using 1,397 single copy orthologous genes, conserved among the 157 strains in the *P. polymyxa* group (**Figure 1B & S4**). The different species and subgroups appear clearly and are robustly supported by Shimodaira-Hasegawa values (Shimodaira & Hasegawa, 1999). The strain identified as species group C serves as outgrouping sample.

Genome completeness levels are predicted to be similarly distributed across strains from the two major species groups. Also, all strains included in the study were analyzed using GTDB-Tk to validate their current taxonomic standing and affiliation to *P. polymyxa* (Chaumeil et al., 2022). The strains have a genome size ranging from 5.15 Mbp for strain YUPP-8 to 6.57 Mbp for strain ISL-58. The lowest GC content is found in strain SC2 with 44.6% and the highest in strain YUPP-8 with 46.6% (**Figure 1C & Table 1**). Strains of species group A have a tendentially smaller genome with lower GC% content than species group B strains.

**Table 1.** Genomic features of *P. polymyxa* strains (n=157). The average genome size in base pairs, number of coding sequences, coding density, GC percentage, number of ribosomal RNAs and number of non-coding RNAs were calculated for each species group and subspecies group.

| Species group | Genome size (bp) | CDSs | Coding density | GC (%) | rRNAs | ncRNAs |
| --- | --- | --- | --- | --- | --- | --- |
| **Group A (*P. polymyxa*)** | **5,813,992** | **5,166** | **86.9** | **45.35** | **20.7** | **6.7** |
| Subgroup A1 | 5,858,337 | 5,237 | 86.2 | 45.13 | 24.4 | 7.0 |
| Subgroup A2 | 5,804,532 | 5,150 | 87.1 | 45.40 | 20.0 | 6.6 |
| **Group B** | **5,861,318** | **5,238** | **86.2** | **45.55** | **18.4** | **7.4** |
| Subgroup B1 | 5,964,228 | 5,283 | 86.1 | 45.57 | 18.9 | 7.5 |
| Subgroup B2 | 5,804,772 | 5,221 | 86.4 | 45.44 | 13.4 | 6.7 |
| Subgroup B3 | 5,758,371 | 5,199 | 86.3 | 45.66 | 18.4 | 7.4 |
| **Group C** | **6,570,682** | **5,795** | **86.1** | **45.80** | **5.0** | **9.0** |

Gene detection was conducted *de novo* on all genomic sequences to homogenize their annotation using Bakta (Schwengers et al., 2021). Then, pangenome analyses were performed using Ppanggolin (Gautreau et al., 2020). For every pangenome analysis, the rarefaction of the total number of genes indicates that there is still strain diversity to be found within the different species as no saturation of gene diversity is reached (**Figure 1D & 1E**). The core genomes of species group A and B represent a high fraction of their total genome with 3,004 and 2,247 CDSs respectively (Supplementary data). Expectedly, species group A shows more conservation as suggested by the ANI levels (**Figure 1A**). The larger strain diversity in species group B is also displayed through the size of its pangenome with 18,203 CDSs compared to 13,889 for species group A (Supplementary data). Another metric depicting the higher genome dissimilarity in species group B is the genomic fluidity of 0.167 compared to 0.12 in species group A (Kislyuk et al., 2011). Finally, unique genes are more abundant on average in species group B with 104 copies per strain compared to 31 in species group A. The persistent genomes of species group A and B comprise 4,129 and 4,014 CDSs respectively or 79.93% and 76.63% of their mean gene number (Supplementary data).

The detected coding sequences were further compared to the PlaBA-db and Annotree databases (Gautam et al., 2022; Mendler et al., 2019; Patz et al., 2021, 2024). In both cases, gene function does segregate the different strains into the previously determined species groups, suggesting different metabolic capacities and different ecologies (**Figure 1F & 1G**).

### The *P. polymyxa* species groups display several distinct metabolic functions

While *P. polymyxa* is generally described as a nitrogen-fixating species, the irregular presence of this trait has been repeatedly observed (Xie et al., 2016; L. Zhou et al., 2020). Indeed, the presence of the *nifHDKENB* gene set is strongly correlated with the genomic background of species group B. The genes responsible for nitrogen fixation can be found in 87.7% of species group B, while they are only present in 5.5% of species group A and strictly restricted to the species subgroup A1 (**Figure 2A**). Several nutrient transport systems were found to be segregating between species group. An operonic structure containing genes putatively involved in polyamine import was found exclusively in strains with species group B background (**Figure 2B**). In species group A, strains are enriched with the *frl* and *lic* operons, respectively responsible for fructoselysine and β-glucoside utilization (**Figure 2C & 2D**). While the *frl* operon is rare in species group B, a modified *lic* operon is common in the species subgroup B1. It contains an additional 6-phospho-β-glucosidase. A 42.9 kb-long genomic region responsible for secondary metabolism was detected to be preferentially present in species group A strains (**Figure 2E**).

**Figure 2.**
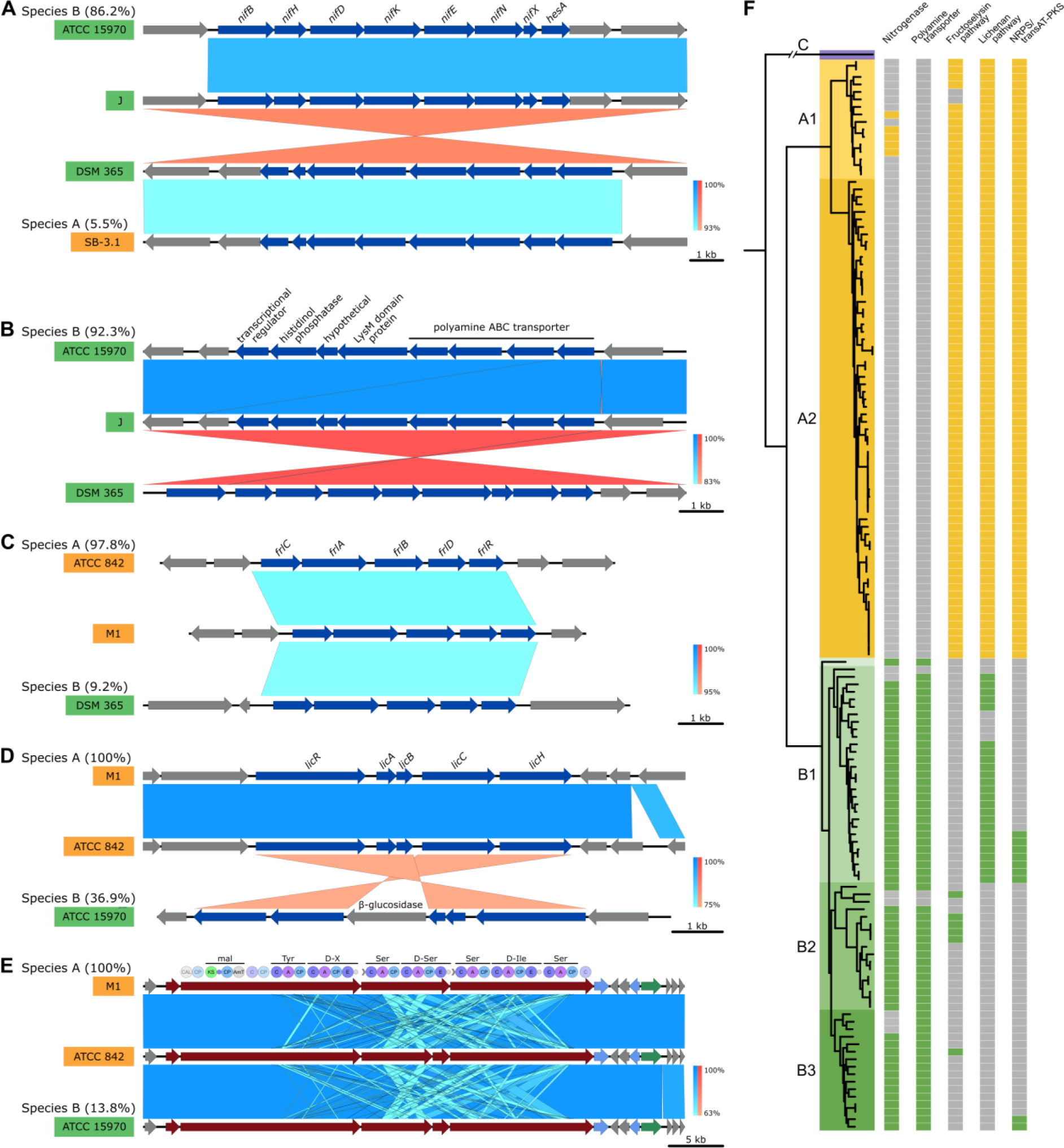
Conserved enrichments of genetic pathways in *P. polymyxa* species groups. The synteny of a non-exhaustive selection of genetic pathways with differential abundances depending on the species group as inferred through Lefse (LDA >1.9, pval<1E-5) has been generated using EasyFig. In each case, the synteny consists of one representative of each species subgroup where the genomic pathway was detected. The total ratio of strains possessing the pathway is given for each species group. Similarity levels were inferred through blastn and are displayed using a blue color scale or a red color scale for inverted sequence identities. For the nitrogenase pathway (A) the polyamine transporter pathway (B) the fructoselysine degradation pathway (C) and the lichenan degradation operon (D) genes with a significant Lefse output (blue) are distinguished from the remaining ones (grey). For the NRPS/transAT-PKS hybrid system (E) genes were colored according to the AntiSmash color code distinguishing biosynthetic (red), transport (blue) and regulatory (green) genes. In addition, the module domain-composition is provided. A detailed description of the presence absence profiles of the different pathways in the *P. polymyxa* strains (F) can be found as supplemental data.

The pathway is predicted to function as a trans-acyltransferace PKS/NRPS hybrid system and to generate a metabolite unknown to the antiSMASH database.

*P. polymyxa* has been repeatedly reported to possess multiple antimicrobial resistances (Eastman et al., 2014; Jeong et al., 2019; Pasari et al., 2019; Storm et al., 1977; L. Zhou et al., 2020). Indeed, using the Comprehensive Antibiotic Resistance Database (CARD) complemented with a similarity search for resistance genes reported in *P. polymyxa* A18 (Pasari et al., 2019), the *P. polymyxa* group displays genes for resistance against penicillin, tellurium, daunorubicin, fosmidomycin, erythromycin, kanamycin, teicoplanin, chloramphenicol, clindamycin, fosfomycin, tetracycline, vancomycin as well as benzalkonium chloride. However, these resistance genes are not all conserved across the different species groups. Species subgroups B1 and B3 are fully devoid of erythromycin resistance genes and only 47% of species group A2 possess them. Resistance against daunorubicin was lost in 73% of species subgroup A2 whereas kanamycin resistance was lost in 69% of species subgroup B3. All other resistance genes were detected in more than 70% of strains from each group (**Figure 3**).

**Figure 3.**
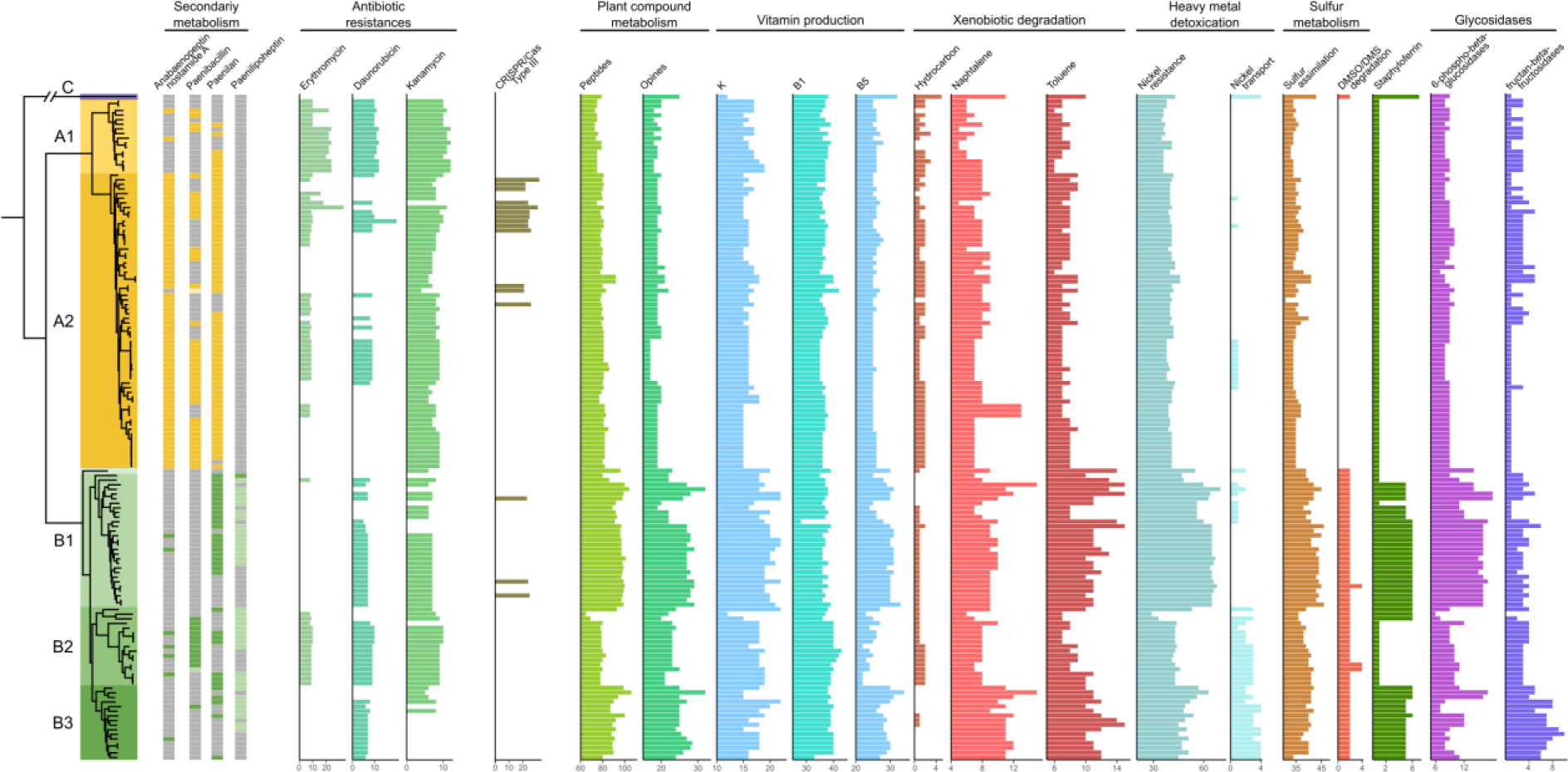
Gene of interest distribution across *P. polymyxa* strains. Genes involved in secondary metabolism were inferred through antiSMASH. Dark-color squares indicate the presence of pathways with >90% similarity. Light-color squares represent incomplete pathways with >40% similarity. Antibiotic resistance genes were detected using the CARD database. The CRISPR/Cas systems were detected using MacSyFinder. All remaining functions were detected with the pgpt-db. The list is not exhaustive, and only pathways with differential distributions between species groups or subgroups are represented.

Genome were screened for the presence of macromolecular systems using MacSyFinder (Abby et al., 2014). The presence of a Tad-pilus and a flagellum is ubiquitous, except for species group C lacking the flagellum coding genes. In accordance to what has been reported in a previous comparative genomics approach, the Gram positive twin-arginine translocation pathway (*tatAC*) was found in all *P. polymyxa* strains (Eastman et al., 2014). All the components for the general secretory pathway *secAYEG* were also detected.

*P. polymyxa* A 18, a gut isolate, was screened for CRISPR/Cas systems putatively conferring resistance against phage invasion which are present in high numbers in the gut environment. Out of the 157 strains screened here, 77 across all species groups except species group C present complete CRISPR/Cas systems. Most of these strains display a Type I system against double stranded DNA (20.8% species group A, 46.2% species group B). However, the Type III system, targeting single strand RNA, is enriched in species subgroup A2 with 15 positive strains compared to only three in species group B (**Figure 3**).

A screening of the secondary metabolism of the 157 strains using antiSMASH revealed interesting patterns for the production of anabaenopeptin NZ857/nostamide A, paenibacillin, paenilan and paenilipoheptin (Blin et al., 2023) (**Figure 3**). The complete metabolic pathways to synthetize fusaricidine were detected in all but five strains. The capacity to synthetize polymyxin is conserved in all but twelve strains, distributed in the two major species groups with no apparent segregating pattern (Supplementary data).

From the pgpt-db we can infer what functional categories contribute most to the segregation between *P. polymyxa* species groups (**Figure 1F**). Gene function linked to plant growth promotion were essentially found to be enriched in species group B compared to species group A (**Figure 3**). Species group B could benefit from an enhanced capacity to utilize plant derived compounds, improving its competitivity in the rhizosphere environment. Especially the species subgroup B2 subgroup is enriched in genes functioning in peptide, opine and beta-glucan utilization. species subgroup B3 displays an enrichment in gens for fructan glycolysis. The direct benefits of species group B for plants can be derived from genes functioning in vitamin production, xenobiotic degradation, nickel detoxication, iron acquisition and sulfur assimilation (**Figure 3**).

Additional functions, directly or indirectly linked to plant growth promotion potential, such as nitrogenase activity, antimicrobial resistance or CRISPR-Cas systems were developed in more detail above.

## Discussion

Comparative genomics analyses were performed on 157 *P. polymyxa* strains, resulting in a composite description of the species. Whole-genome alignments and phylogenomic reconstructions strongly suggest the presence of two dominant species groups that do not share a monophyletic position on the *Paenibacillus* genus tree. Ninety-one strains share genomic and functional similarities with the type strain ATCC 842 of *P. polymyxa*, while 65 strains show strong evidence for the existence of a separate species. Further based on genomic and functional criteria, several subgroups can be delimited within each *P. polymyxa* species group.

The genomic diversity of *P. polymyxa* strains has been a source of confusion in the past. In 2001, an olive-mill wastewater isolate, strain KACC 10925, was compared to *P. polymyxa* CECT 153 and resulted in significantly low DNA-DNA relatedness, prompting the authors to propose *Paenibacillus jamilae* as a new species (Aguilera et al., 2001). Almost 20 years later, it was argued that when using the type strain *P. polymyxa* ATCC 842, the comparison with KACC 10925 yields sufficiently high similarity to be classified as the same species, relegating *P. jamilae* as a heterotypic synonym of *P. polymyxa* (Kwak et al., 2020). This confusion arose from the heterogeneity of strains classified as *P. polymya*, which we described here. Strain CECT 153 is part of the species B group, which is indeed genomically distant from the type strain group, including KACC 10925.

We have put the focus on the plant interactive traits which are often reported for *P. polymyxa* (L. Zhou et al., 2020). As it is a promising plant growth promotion agent, it is intriguing to verify how the different species groups differ in their genetic baggage dedicated for plant interaction. A major recognition, is that the inconsistent distribution of nitrogen fixation genes, repeatedly described in the literature, is heavily correlated with the genetic background of the strain (Xie et al., 2016; L. Zhou et al., 2020). This feature is only rarely found in strains with the genetic background of the *P. polymyxa* type strain, and largely enriched in the newly separated species. This species group shows further enrichment in genes directly or indirectly involved in plant growth promotion, as well as plant substrate usage.

The pangenomic analysis revealed presence/absence patterns for further complete metabolic pathways, correlating with the different genetic backgrounds of the *P. polymyxa* species groups. The putative polyamine transporter detected exclusively in the species group B background could also have an impact on plant interaction. *Pseudomonas* sp. WCS365 uses the polyamine putrescine as a signaling molecule with its host during colonization of *Arabidopsis thaliana* (Liu et al., 2018). The implication of putrescine-related pathways in plant colonization was also underlined for *Burkholderia* species interacting with rice (Wallner et al., 2022, 2023). Polyamines further play a role in biofilm formation in different bacterial models (Karatan & Michael, 2013).

The *P. polymyxa* type strain species group and the species group B group display a strong conservation in secondary metabolism. Both species groups retain the capacity to synthetize polymyxin, often described as a last resort antimicrobial against multidrug-resistant pathogens (Zavascki et al., 2007). They also have the capacity to produce fusaricidin a potent antimicrobial against Gram-positive bacteria and phytopathogenic fungi (Li & Chen, 2019; Yu et al., 2012). However, a predicted trans-acyltransferase PKS/NRPS system is conserved in every strain of species group A, while it is only present in few species group B strains. It is predicted to be composed of seven amino acids but displays no similarity to any synthesis pathway in secondary metabolite databases. Its strong conservation level within the *P. polymyxa* type strain group make it an interesting target for future research.

The *<u>lic</u>* operon was studied in *Bacillus subtilis* and allows the bacterium to use oligomeric β-glucosides as sole carbon source (Tobisch et al., 1999). In *Clostridiodes difficile* it enables the utilization of cellobiose (Hasan et al., 2021). Cellobiose is a product of cellulose degradation, but it can also be found in maize grains and is present in maize root where it participates in immune responses (Saravanakumar et al., 2016). The presence of the *lic* operon could benefit organisms associating with plants producing this sugar or with cellulose-degrading microorganisms, scavenging the breakdown compounds. Interestingly, the species subgroup B1 subgroup possesses a *lic* operon containing an additional gene identifies as a 6-phospho-β-glucosidase. This function is also found in the cellobiose-utilization operon of *Escherichia coli* (Verma & Mahadevan, 2012). While in *E. coli* this function is essential for the utilization of cellobiose, a homologous gene is present outside of the operon in strains of species group A. The *frl* operon is almost completely absent in species group B but highly conserved in species group A. It enables the use of fructoselysine, a stable glycation product of glucose and lysine formed nonenzymatically (Wiame et al., 2002).

Both cellobiose and fructoselysine are degradation products and found in animal digestive systems where they are used by intestinal bacteria. All *P. polymyxa* strains originating from animal associated environments belong to the genomic background where *frl* and *lic* operons are enriched. This species group is also enriched in antimicrobial resistance genes as well as in CRISPR-Cas systems which could benefit strains in environment with increased viral pressure such as the gut environment.

In conclusion, our approach of combining multiple bioinformatics analyses to compare 157 genomes of *P. polymyxa* at the genomic and functional level revealed the existence of two species groups, as 65 strains show significant genomic divergences and specific functional traits compared to the 91 strains showing similarities to the type strain *P. polymyxa* ATCC 842.

## Methods

### Genomic data

Genome sequences belonging to *P. polymyxa* were downloaded from NCBI’s RefSeq database. Additional genome sequences referenced as *P. polymyxa* on GTDB were downloaded from NCBI’s GenBank database (Supplemental data). The homogenous affiliation to *P. polymyxa* was verified using the GTDB-Tk v2.1.1 classify workflow (Chaumeil et al., 2022; Eddy, 2011; Hyatt et al., 2010; Jain et al., 2018; Matsen et al., 2010; Ondov et al., 2016; Price et al., 2010). Genome completeness levels were assessed using the checkM v1.2.2 lineage workflow (Parks et al., 2015). All assemblies were annotated *de novo* using Bakta v1.9.2 with baktaDB5.1 to standardize protein annotation between strains (Schwengers et al., 2021).

### Average nucleotide identity analyses

ANI calculations were conducted with Pyani v0.2.12 (Pritchard et al., 2016) using either BlastN (ANIb) or MUMmer (ANIm) supported alignments (Altschul et al., 1990; Kurtz et al., 2004). Standard threshold for species delimitation were considered i.e., 95% identity with 70% sequence coverage. ANIb was used to compare the nucleotide sequences from the 157 strains affiliated to *P. polymyxa* and ANIm to compare the type strains of 93 *Paenibacillus* species.

### Phylogenetic reconstructions

Detection of conserved gene orthologues was conducted separately on the complete set of 157 genomes affiliated to *P. polymyxa* and to the same set with five additional *P. ottowii* genomes using OrthoFinder v2.5.5 (Emms & Kelly, 2019). The aligned amino acid sequences of single copy orthologous genes were used for approximately-maximum-likelihood phylogenetic tree inference using FastTree v2.1.11 (Price et al., 2010). The JTT model of amino acid evolution with CAT approximation and a Gamma20-based likelihood branch rescaling was used. Local support values are provided through the Shimodaira-Hasegawa test.

### Pangenome analyses

Gene conservation across strains was calculated using PPanGGOLiN v2.0.2 on bakta-generated .gbff files (Gautreau et al., 2020). Separate analyses were run for the total of 157 *P. polymyxa* affiliated strains, strains belonging to species group A and strains belonging to species group B.

### Functional screening

Protein sequences were annotated and analyzed with the PGPT_BASE_nr_Aug2021n_ul_1 database from PLaBAse and the AnnoTree Bacteria database (version of 25 August 2020) using the DIAMOND + MEGAN pipeline with DIAMOND v2.1.8 and MEGAN v6.24.23 (Bağcı et al., 2021; Gautam et al., 2022; Mendler et al., 2019; Patz et al., 2024). Functional cluster analyses were generated using PCoA based on Bray-Curtis distances.

GenBank format annotation and sequence files were used as input to screen for secondary metabolite synthesis pathways using antiSMASH v.6.1.1 (Blin et al., 2021). Only contigs with a length of 500 nt or more were considered for the analysis consisting of a whole-genome HMMer scan, active site finder and comparison to antiSMASH’s database of known cluster.

For antimicrobial resistance screening, the CARD database v3.2.9 was used with the resistance gene identifier tool v 6.0.3 on protein data files using the DIAMOND alignment tool (Alcock et al., 2023). Additionally, resistance genes that were not detected by alignment to the CARD databases but mentioned in (Pasari et al., 2019) i.e., penicillin, tellurium, daunorubicin, fosmidomycin, erythromycin, kanamycin, teicoplanin and chloramphenicol resistance genes, were screened manually using the reference sequences provided by the authors.

Macromolecular systems and CRISPRS-Cas systems were detected using MacSyFinder v2.1.3 with TXSScan v1.1.3 and CasFinder v3.1.0 models (Abby et al., 2014, 2016; Néron et al., 2023).

### Synteny analyses

Genomic regions of interest were detected by applying a differential abundance analysis on the pangenome output using LEfSe v1.1.2 (Segata et al., 2011). Species groups were used as class, Species subgroups were used as subclass and individual IDs were used as subject for the analysis. Synteny comparison figures were drawn using Easyfig v2.2.5 (Sullivan et al., 2011). For every region of interest, one representative of each species subgroup in which the region is fully or partially present is depicted. The similarity scores were inferred through blastn (Altschul et al., 1990).

## Declarations

### Availability of data

All genomes used in this study are available on NCBI’s GenBank database (https://www.ncbi.nlm.nih.gov/genbank/).

### Author’s contribution

AW analyzed and interpreted the genomic data. LA, OM, PR and SC made substantial contribution during study conception and manuscript improvements. All authors read and approved the final manuscript.

## Supplemental data

**Figure S1.**
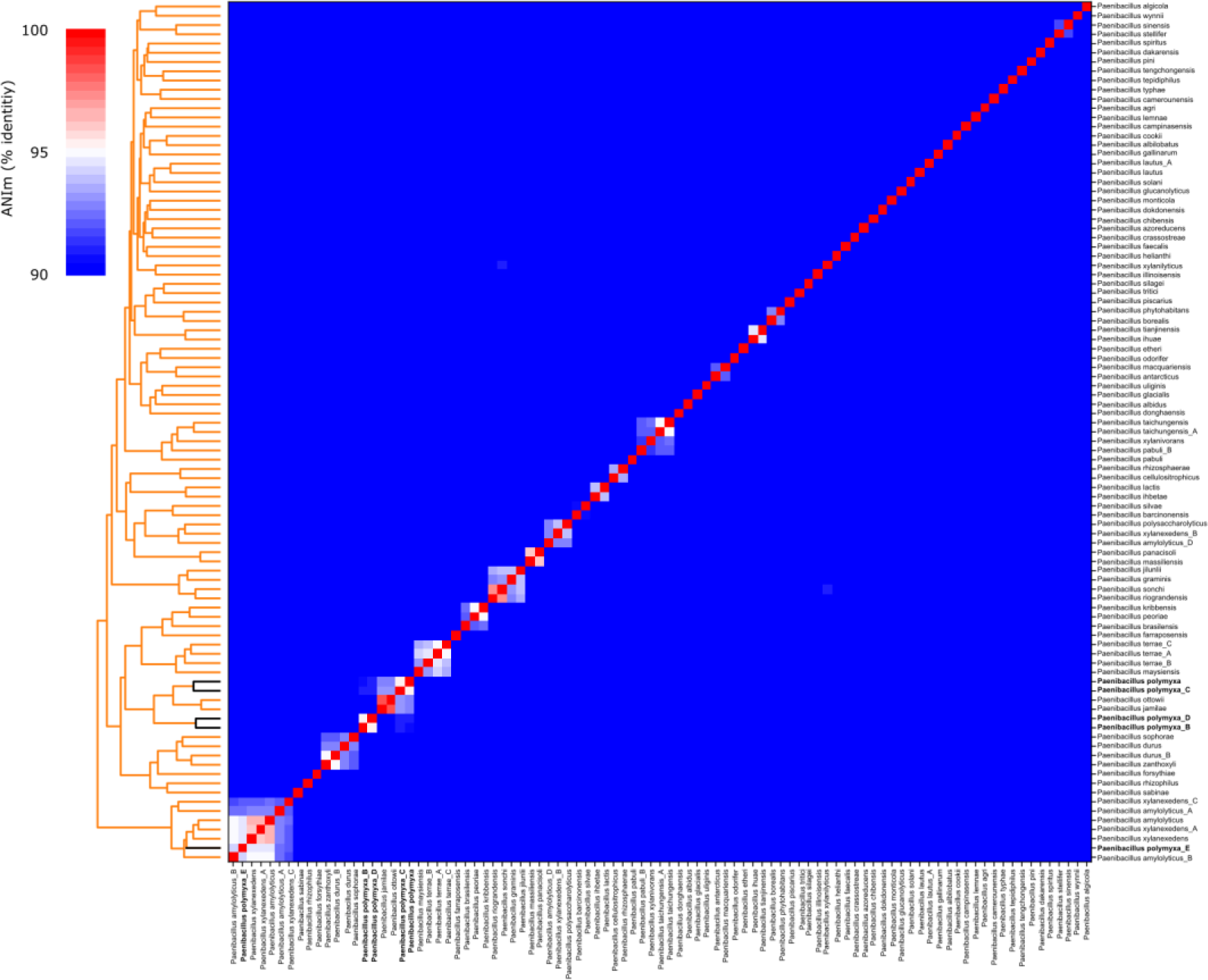
Average nucleotide identity results for 93 *Paenibacillus* species representative strains based on mummer comparisons. The representative strains of GTDB-validated *Paenibacillus* species were selected for this analysis. Strains of the different *P. polymyxa* groups are indicated in bold and black bars on the dendrogram.

**Figure S2.**
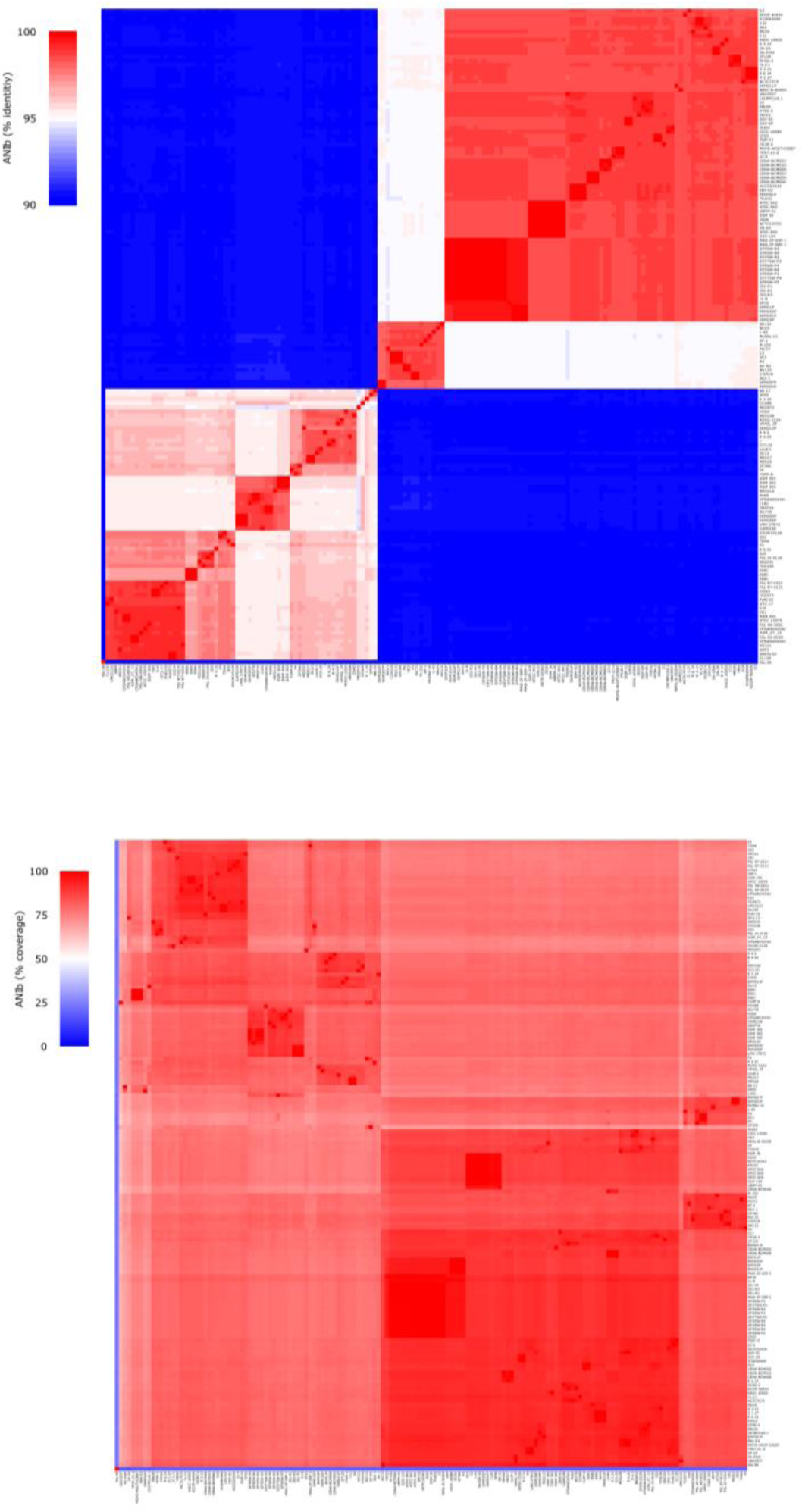
Average nucleotide identity results for 157 *P. polymyxa* strains based on blast comparisons. (A) Percentage identity and (B) percentage alignment coverage.

**Figure S3.**
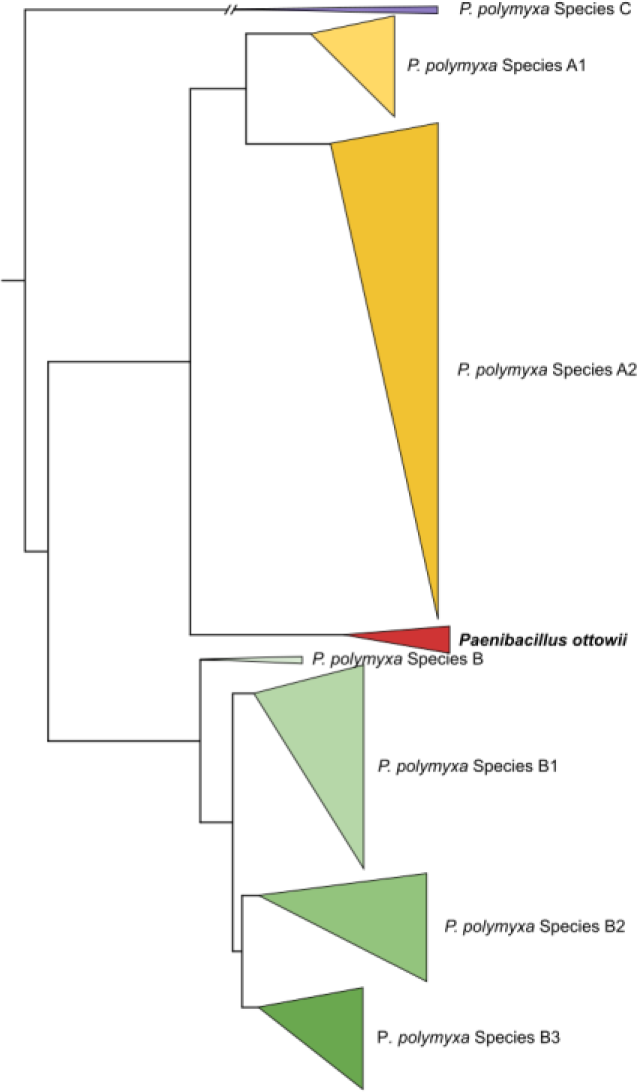
Phylogenetic reconstruction *P. polymyxa* species groups and *P. ottowii*. Single copy orthologous protein sequences were detected and aligned for 157 *P. polymyxa* genomes and 5 *P. ottowii* genomes using Orthofinder resulting in an alignment of 1388 sequences for a total of 403,770 amino acids. The tree was build using FastTree with default parameters.

